# Elevated mutation near crossovers inhibits the evolution of recombination

**DOI:** 10.1101/2025.11.01.685904

**Authors:** Bret A. Payseur, Sarah P. Otto

**Affiliations:** Laboratory of Genetics, University of Wisconsin-Madison, Madison, Wisconsin, U. S. A.; Department of Zoology and Biodiversity Research Centre, University of British Columbia, Vancouver, British Columbia, Canada

**Keywords:** recombination, deleterious mutation, epistasis, double-strand break, DNA repair

## Abstract

Recombination diversifies offspring genomes and helps ensure chromosome segregation during meiosis. Mutation rates are elevated near crossovers due to the induction of double-strand breaks and their imperfect repair, a byproduct of recombination typically ignored by theory designed to explain its evolution. To examine the evolutionary role of recombination-associated mutation, we analyze a population genetic model in which a modifier locus controls both the rate of recombination between two loci experiencing viability selection and the rate of mutation at those loci. Analytical and numerical results demonstrate that the advantage of recombination conferred by its capacity to remove epistatic, deleterious mutations is overcome by the selective cost of even small increases in the mutation rate. Simulations of finite populations show that recombination-associated mutation also decreases the probability of fixation of modifier alleles in the absence of epistasis. By incorporating the mutagenic effects of recombination, our analysis extends a rich body of theory to incorporate a molecular feature inherent in the formation of crossovers. Our findings suggest that higher recombination rate evolves by altering steps in the crossover pathway that are less likely to inflict mutational damage.

## Introduction

During meiosis, homologous chromosomes exchange DNA via crossovers, creating new combinations of genetic variants. Recombination improves the efficiency of natural selection, shapes genomic patterns of variation, and (in many species) helps to ensure that chromosomes segregate properly (Fisher 1930; Muller 1932; Hill and Robertson 1966; Felsenstein 1974; Feldman et al. 1980; Maynard Smith and Haigh 1974; Begun and Aquadro 1992; Charlesworth et al. 1993; Barton 1995; Feldman et al. 1996; Otto and Barton 1997; Otto and Feldman 1997; Hassold and Hunt 2001; Page and Hawley 2003; Cutter and Payseur 2013; Charlesworth and Jensen 2021). Therefore, explaining heterogeneity in recombination rate observed along the genome, among individuals, and between species (McVean et al. 2004; Coop and Przeworski 2007; Smukowski and Noor 2011; Stapley et al. 2017; Haenel et al. 2018; Henderson and Bomblies 2021; Johnston 2024) is a goal with broad evolutionary significance.

Population genetics theory has established conditions that favor the evolution of recombination. One powerful approach is to model the fate of alleles at a modifier locus that controls the rate at which other loci recombine (Nei 1967; Feldman 1972). In large populations at equilibrium, recombination-reducing modifiers become associated with beneficial combinations of alleles and spread, leading to evolutionary decreases in recombination (Feldman et al. 1980; Feldman and Liberman 1986; Barton 1995). Multiple scenarios can lead to the evolution of higher recombination rate. In finite populations, genetic drift interacts with natural selection to generate linkage disequilibrium between beneficial and deleterious alleles; this “selective interference” confers an advantage to recombination-increasing alleles that reduce these associations and increase the efficacy of selection (Hill and Robertson 1966; Hill and Robertson 1968; Peck 1994; Otto and Barton 2001; Barton and Otto 2005; Keightley and Otto 2006).

Temporal and spatial fluctuations in selection can favor recombination-increasing modifiers if combinations of alleles that are advantageous in one environment become detrimental (Barton 1995; Otto and Michalakis 1998; Barton and Charlesworth 1998; Peters and Lively 1999; Lenormand and Otto 2000; Gandon and Otto 2007). When deleterious alleles at two loci reduce fitness more than expected from their independent effects (synergistic or negative epistasis), linkage disequilibria develop that reduce genetic variance, in which case, higher recombination can be favored if a modifier that elevates recombination increases the efficiency of selection enough relative to the recombination load that it induces (Feldman et al. 1980; Otto and Feldman 1997; Barton and Charlesworth 1998).

Although mutation is often a key ingredient in models of the evolution of recombination, recombination and mutation are routinely treated as independent processes. Consideration of the molecular and cellular steps that lead to the formation of crossovers challenges this assumption. Recombination requires the programmed generation of DNA damage in the form of double-strand breaks (DSBs), a minority of which are resolved as crossovers between homologous chromosomes (Keeney et al. 1997; Mahadevaiah et al. 2001; Martini et al. 2011; Cole et al. 2012; Yokoo et al. 2012; Hunter 2015; Varas et al. 2015; Gray and Cohen 2016). The repair of DSBs associated with recombination is prone to error (Macaisne et al. 2018; Rodgers and McVey 2016; Hanscom and McVey 2020), causing recombination to be mutagenic. The most direct evidence for this claim comes from experiments in budding yeast. After Magni and Von Borstel (1962) used the frequency of reversion of auxotrophic alleles to conclude that the mutation rate is higher during meiosis than mitosis, Magni (1963) showed that a large excess of meiotic revertant cells was associated with a crossover. Mitotic budding yeast cells with DSBs experimentally induced near revertible alleles have a 100-fold higher reversion rate than cells without induced DSBs (Strathern et al. 1995). Using a forward mutation reporter, Rattray et al. (2015) found an elevated mutation rate at a meiotic recombination hotspot (compared to a coldspot). The higher mutation rate is dependent on the protein that generates DSBs during meiosis (*SPO11*), tying mutagenic recombination to the processing of DSBs (Rattray et al. 2015).

Comparing the locations of crossovers and *de novo* mutations observed in human pedigrees provides additional evidence for elevated mutation with recombination. Halldorsson et al. (2019) estimated that within 1kb of crossovers, maternal and paternal mutation rates are 58.4x and 41.5x their respective genome-wide rates. Examining the overlap between mutations and recombination hotspots (inferred from binding of the recombination intermediate protein *DMC1*), Hinch et al. (2023) suggested that 1 in 4 sperm and 1 in 12 eggs harbors a new mutation specifically due to DSB repair. Positive correlations between recombination rate and among-species sequence divergence extend support for recombination-associated mutation to non-human primates (Rouselle et al. 2019), flycatchers (Bolivar et al. 2016), fowls (Rouselle et al. 2019), and *Drosophila* (Begun et al. 2007; Kulathinal et al. 2008), though this evidence is less direct than that found in experimental and pedigree-based studies.

Current models of the evolution of recombination ignore the possibility of recombination-associated mutation. To address this gap in the field, we extend population genetic theory to simultaneously consider changes in the recombination rate and changes in the mutation rate. We discover that modest and realistic increases in mutation rate near crossovers impose a formidable barrier to evolving more recombination.

## Model and Results

Our goal is to determine how recombination-associated mutation affects the evolution of recombination driven by selection against deleterious variants. We build on the modeling framework established by Feldman et al. (1980) and Otto and Feldman (1997), who demonstrated that weak, synergistic epistasis against deleterious mutations favors the evolution of higher recombination. Following this analytical treatment, we conduct stochastic simulations that incorporate recombination-associated mutation into a model of selective interference (Keightley and Otto 2006).

We assume a haploid population that is large enough that genetic drift can be ignored.

We analyze a three-locus modifier model, following changes in haplotype frequencies from one generation to the next, as illustrated in **Figure 1**. We census at the adult haploid stage, following which random mating creates diploids, which then undergo meiosis. During meiosis, the modifier locus (*M*) controls the rates of mutation and recombination at two loci that affect viability (*A* and *B*), arranged in the following order on the chromosome: *M-A-B*. The *A* and *B* alleles each experience recurrent mutation to deleterious alleles (*a* and *b*, respectively) at rate *u*_*MM*_ in haplotypes homozygous for the resident allele at the modifier locus (*MM*), *u*_*Mm*_ = *u*_*MM*_(1 + *d*) in haplotypes that contain both resident and new modifier alleles (*Mm*), and *u*_*mm*_ = *u*_*MM*_(1 + 2*d*) in haplotypes homozygous for the new modifier allele (*mm*). Reverse mutations and mutations at the modifier locus are ignored. The *d* parameter captures the proportional change in the mutation rate caused by the new modifier allele. The modifier locus recombines with the first viability locus (*A*) at rate *R* (only relevant in *Mm* heterozygotes). The rate of recombination between viability loci *A* and *B* is determined by the genotype at the modifier locus: *MM* individuals recombine at rate *r*_*MM*_, *Mm* individuals recombine at rate *r*_*Mm*_, and *mm* individuals recombine at rate *r*_*mm*_. Here, we assume that elevated levels of DSBs induce higher mutation rates, whether or not a crossover results in the region (similar results are obtained when elevated mutation is coupled with recombination). Following meiosis, selection acts on haploid viability according to the haplotype at loci *A* and *B*. The viabilities (fitnesses) of *AB, Ab, aB*, and *ab* individuals are 1, 1 − *s*, 1 − *s*, and (1 − *s*)^2^ + *ϵ*, respectively, where *s* denotes the loss of viability caused by selection and *ϵ* denotes epistasis between the two loci. When *ϵ* is negative, the fitness of the *ab* combination is less than expected from the product of the fitnesses of the constituent alleles (synergistic or negative epistasis). With two alleles at each of three loci, there are eight possible haplotypes {*MAB, MAb, MaB, Mab, mAB, mAb, maB, mab*} whose frequencies among adult haploids are {*x*_1_, *x*_2_, *x*_3_, *x*_4_, *x*_5_, *x*_6_, *x*_7_, *x*_8_}. Parameter abbreviations are summarized in **Table 1**.

**Table 1.** Abbreviations for model parameters.

| Abbreviation | Parameter |
| --- | --- |
| $R$ | Rate of recombination between modifier locus ($M$) and first viability locus ($A$) |
| $r_{MM}, r_{Mm}, r_{mm}$ | Rates of recombination between first viability locus ($A$) and second viability locus ($B$) for the $MM$, $Mm$, and $mm$ genotypes (respectively) at the modifier locus |
| $u_{MM}, u_{Mm}, u_{mm}$ | Rates of mutation at the two viability loci ($A$ and $B$) for the $MM$, $Mm$, and $mm$ genotypes (respectively) at the modifier locus |
| $d$ | Proportional change in mutation rate conferred by the $m$ allele at the modifier locus |
| $W_1$ | Fitness of haplotype with one deleterious allele ($Ab$ and $aB$) |
| $W_2$ | Fitness of haplotype with two deleterious alleles ($ab$) |
| $s$ | Strength of selection against deleterious alleles ($a$ and $b$) |
| $\epsilon$ | Strength of epistasis; difference in fitness of haplotype with two deleterious alleles ($ab$) from that expected if fitness is multiplicative |

**Figure 1.**
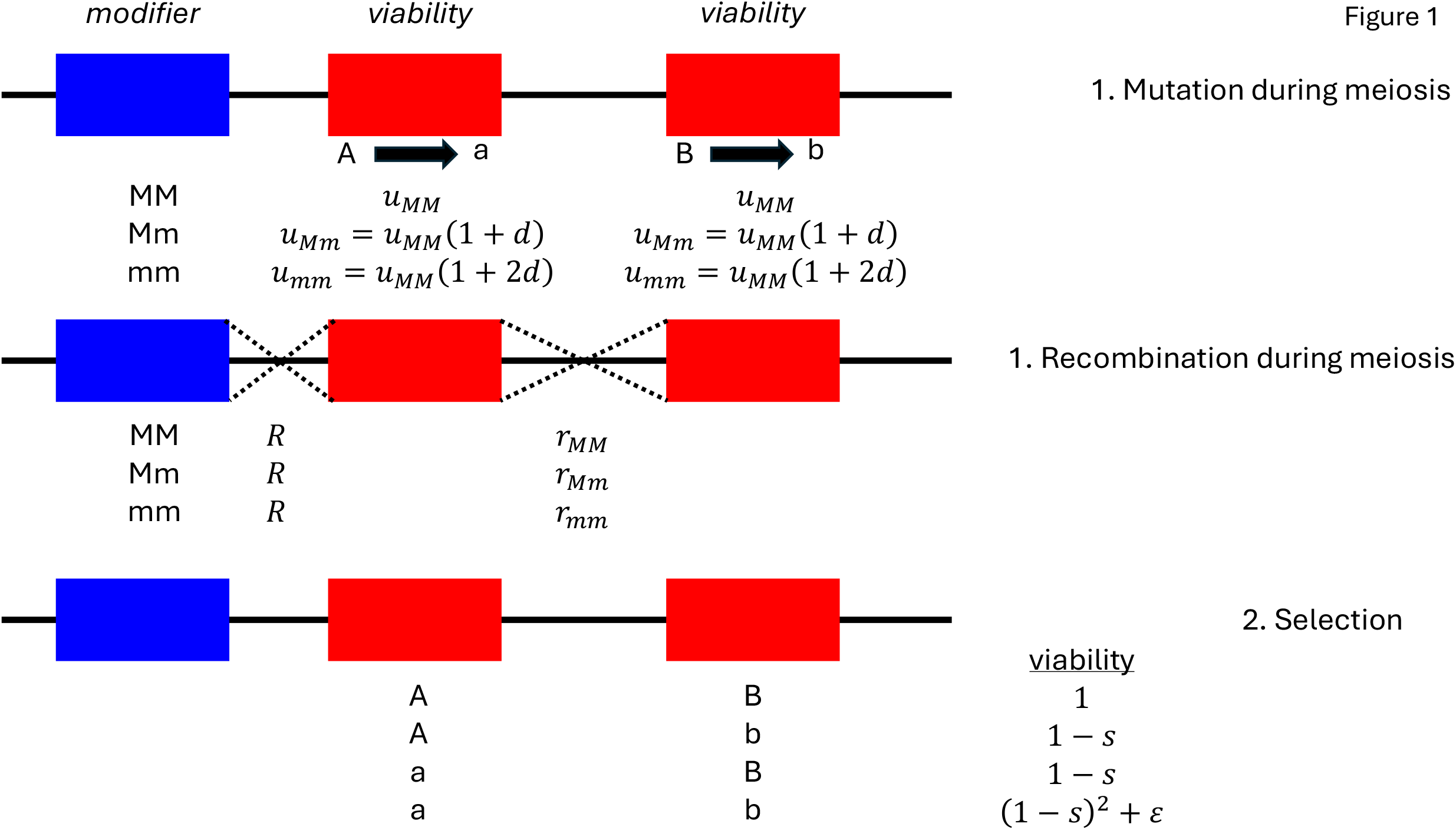
Illustration of the model. A modifier locus (shown in blue) controls rates of recombination and mutation at two loci (red) that undergo recurrent mutation to deleterious alleles. Parameter notation is described in Table 1.

We construct a series of eight recursion equations that describe changes in the eight haplotype frequencies from one generation to the next (Appendix). We seek to determine the conditions under which a modifier allele, *m*, that alters both the recombination rate and the mutation rate, will spread through the population. When the resident allele (*M*) is fixed and selection is strong relative to mutation, the system of recursions for the four haplotypes containing *M* {*x*_1_, *x*_2_, *x*_3_, *x*_4_} equilibrates at a balance between mutation, recombination, and selection at the viability loci (*A* and *B*). We use the frequency of deleterious alleles (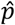) and linkage disequilibrium between alleles (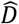) at the two viability loci to describe the polymorphic equilibrium. Assuming epistasis is weak, 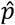 and 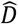 can be approximated to order *ϵ* (Appendix) as:

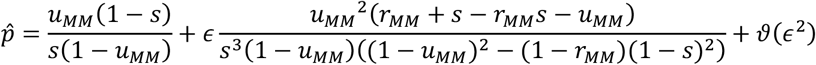

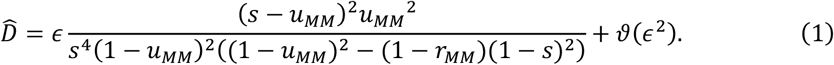

[Note: The denominators in this expression contained a typo in Otto and Feldman (1997), with the mutation rate dropped from (1 − *u*_*MM*_)^2^ − (1 − *r*_*MM*_)(1 − *s*)^2^.] The first part of the expression for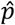 reduces to the familiar ^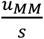^ (Haldane 1937) when *u*_MM_ is small, *s* is small, and *u*_*MM*_ is smaller than *s*.

To evaluate the stability of this resident equilibrium to invasion of the modifier allele, *m*, that affects both recombination and mutation, we construct the stability matrix containing linearized recursions for the frequencies of the four haplotypes that harbor *m* {*x*_5_, *x*_6_, *x*_7_, *x*_8_} (Appendix). The leading eigenvalue of this matrix (*λ*_*L*_) evaluated at the resident equilibrium (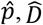) determines the fate of the new modifier allele: invasion occurs when *λ*_*L*_ > 1, whereas invasion is resisted when *λ*_*L*_ < 1. The expression for *λ*_*L*_ for this stability matrix is long and difficult to interpret. To make progress, we assume that epistasis (*ϵ*) and the mutation effect of the modifier allele (*d*) are similarly weak (of order *ϵ*). Then, *λ*_*L*_can be approximated to leading order (Appendix) as:

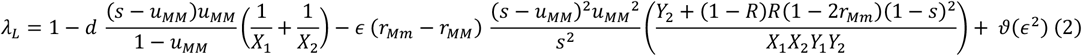

where each fraction is positive under the assumption that mutation is weak relative to selection and the *X*’s and *Y*’s are positive functions introduced to simplify the presentation. Specifically, *X*_1_ is *X*[*R*], *X*_2_ is *X*[*R*(1 − *r*_*Mm*_) + *r*_*Mm*_(1 − *R*)], *Y*_1_ is *Y*[*r*_*MM*_], and *Y*_2_ is *Y*[*R* + *r*_*Mm*_ − *Rr*_*Mm*_], where:

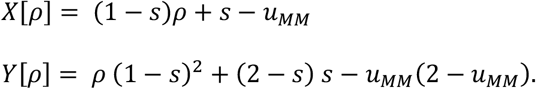

The signs of the two main parts of expression (2) thus determine the fate of a new modifier allele. The sign of the term involving *ϵ* depends on the difference in recombination frequency between modifier genotypes (*r*_*Mm*_ − *r*_*MM*_). When recombination increases with a new modifier allele (*r*_*Mm*_ > *r*_*MM*_) and *ϵ* is weak and negative, the last term is positive, pushing in the direction of invasion (Otto and Feldman 1997; who also show that the *ϵ*-term disfavors increased recombination when epistasis is strong, unless the modifier is very tightly linked). By contrast, the sign of the term involving *d* is determined by the sign of *d* (this term does not depend on the difference in recombination frequency between modifier genotypes). When the new modifier allele increases the mutation rate (*d* > 0), the *d*-term is negative, bolstering the stability of the resident equilibrium. When the new modifier allele decreases the mutation rate (*d* < 0), the *d*-term is positive, favoring invasion. Importantly, the relative magnitudes of these two terms differ, with the *d*-term proportional to *u*_*MM*_ and the *ϵ*-term generally much smaller and proportional to *u*_*MM*_^2^. This difference is more easily seen by assuming weak selection (order *ϵ*) and even weaker mutation (order *ϵ*^2^), in which case the *d*-term reduces to

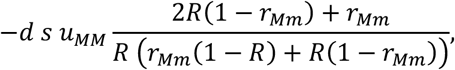

while the *ϵ*-term reduces to

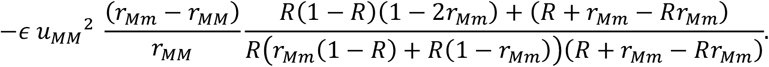

For biologically reasonable values of *d* and *ϵ*, the *d*-term will be larger in magnitude than the *ϵ*-term. Consequently, the fate of the new modifier allele is determined disproportionately by its effect on mutation. A modifier allele that increases both the recombination rate and the mutation rate is not expected to spread through the population, unless DSBs and the mutations they induce are disproportionately targeted to non-functional sites (see Discussion).

To understand how the effects of recombination-associated mutation scale with the number of loci experiencing viability selection, we develop a simpler approximation for the *d*-term of *λ*_*L*_ in (2) and compare it to results from a two-locus model of a mutation rate modifier (Appendix). Our findings indicate that the *d*-term scales nearly linearly with the number of viability loci (Appendix).

The analytical results presented above offer general insights into the evolution of recombination in the realm where epistasis is weak and changes in mutation rate are small. To explore the fate of the new modifier allele under a wider array of conditions, we solve numerically for the resident equilibrium (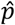 and 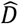) and then for *λ*_*L*_ of the stability matrix under a variety of parameter values.

Figure 2. shows the rate of spread (measured by *λ*_*L*_ − 1) of a modifier allele that increases both recombination and mutation. Numerical results confirm that a modifier allele that increases recombination alone spreads through the population when epistasis is negative and not too strong (**Figure 2A**) (Otto and Feldman 1997). However, if the allele also elevates mutation, it is no longer favored (**Figure 2A**). This disparity persists across a range of selection coefficients (**Figure 2B**). Whereas an allele that increases recombination alone is more likely to spread when the modifier locus is closely linked to the first viability locus (Otto and Feldman 1997), an allele that also elevates mutation is *less* likely to spread with close linkage (**Figure 2C**). Although an allele that increases both recombination and mutation is still favored when recombination levels are initially low, this advantage quickly disappears as the resident recombination rate grows (**Figure 2D**). An allele must expand the recombination rate by a substantial amount to overcome the cost of the small increase in mutation rate it causes (**Figure 2E**). Finally, the spread of an allele that modulates both recombination and mutation is inhibited across a spectrum of resident mutation rates (**Figure 2F**). Overall, numerical results are consistent with analytical results in showing that the advantage of increased recombination in the face of synergistic epistasis among deleterious mutations is eliminated or substantially diminished by the mutational cost inherent in the formation of crossovers.

**Figure 2.**
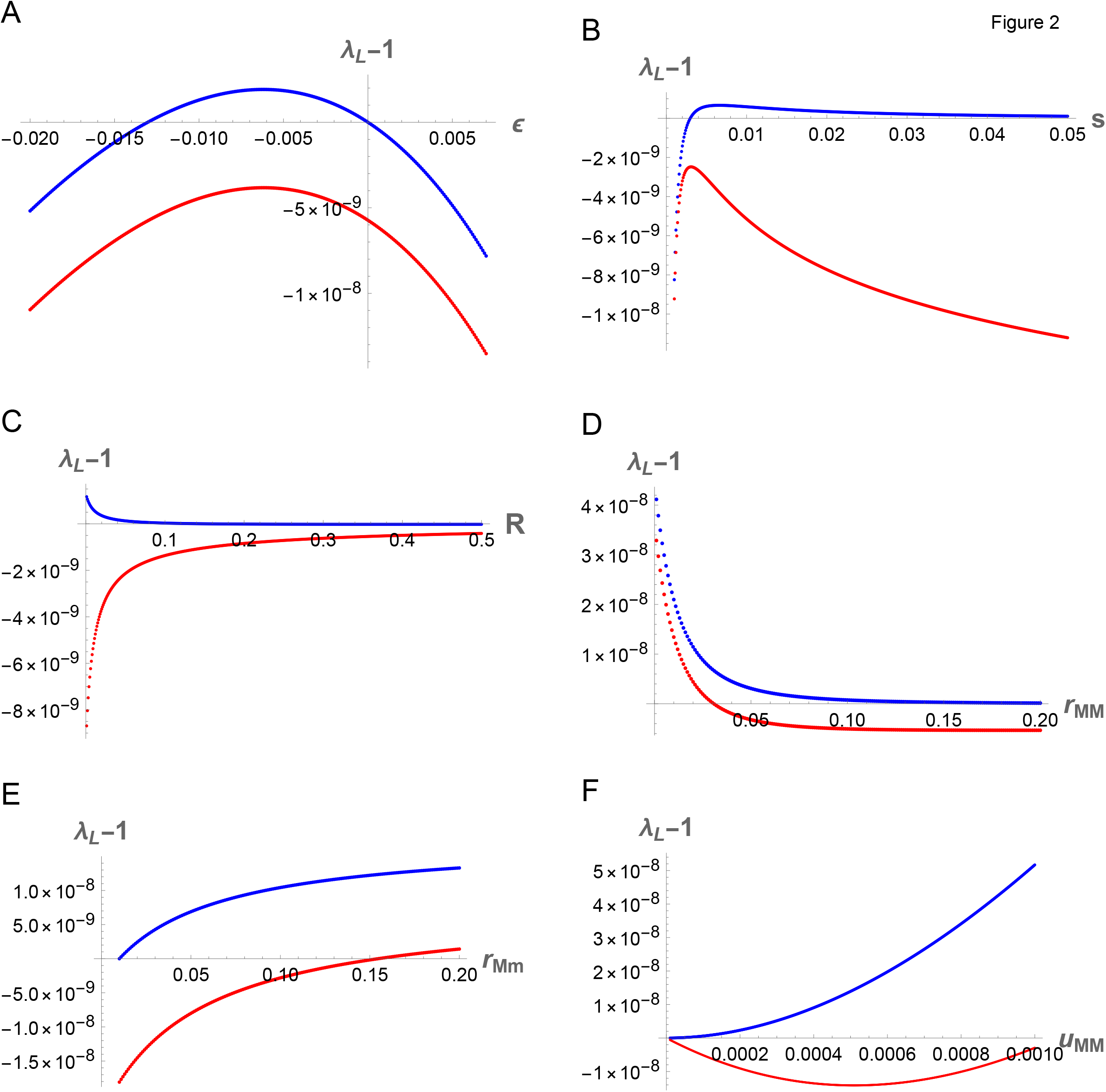
The chances of invasion for a modifier allele that controls recombination rate and mutation rate. Panels A-E show the leading eigenvalue of the stability matrix (*λ*_*L*_) minus 1 for two conditions: (i) the recombination-modifying allele does not affect the mutation rate (*u*_*MM*_ = 0.0001 and *d* = 0; blue), and (ii) the recombination-modifying allele increases the mutation rate by a small factor of 0.01% (*u*_*MM*_ = 0.0001, *u*_*Mm*_ = 0.00010001 (*d* = 0.0001); red). Panel F is similar but allows *u*_*MM*_ to vary. In all cases, invasion occurs when *λ*_*L*_ − 1 is positive. A. Effect of epistasis (*ϵ*). *R* = 0.01, *r*_*MM*_ = 0.10, *r*_*Mm*_ = 0.12, *s* = 0.01. B. Effect of selection (*s*). *R* = 0.01, *r*_*MM*_ = 0.10, *r*_*Mm*_ = 0.12, *ϵ* = -0.001. C. Effect of recombination rate between the modifier locus and the first viability locus (*R*). *r*_*MM*_ = 0.10, *r*_*Mm*_ = 0.12, *ϵ* = -0.001, *s* = 0.01. D. Effect of initial recombination between the two viability loci (*r*_*MM*_). *R* = 0.01, *r*_*Mm*_ = *r*_*MM*_ + 0.01, *ϵ* = -0.01, *s* = 0.01. E. Effect of increase in recombination rate between viability loci (*r*_*Mm*_). *R* = 0.01, *r*_*MM*_ =0.01, *ϵ* = -0.01, *s* = 0.05. F. Effect of baseline mutation rate (*u*_*MM*_). *R* = 0.01, *r*_*MM*_ = 0.10, *r*_*Mm*_ = 0.12, *ϵ* = -0.001, *s* = 0.01.

Synergistic epistasis is not the only evolutionary force capable of generating the linkage disequilibrium that favors evolutionary increases in recombination. Selective interference, caused by the accumulation of beneficial alleles alongside deleterious alleles following a period of selection and genetic drift, also favors the evolution of recombination (Otto and Barton 2001; Barton and Otto 2005; Roze and Barton 2006). Keightley and Otto (2006) used simulations to demonstrate that selective interference among deleterious mutations in finite populations favors the spread of modifier alleles that increase recombination in a model with recurrent deleterious mutations, even in the absence of epistasis. To explore whether recombination-associated mutation inhibits the evolution of recombination in this scenario as well, we conduct individual-based simulations. We modify C code originally written by Peter Keightley (Keightley and Otto 2006) to include recombination-induced mutational effects. We track a haploid population with *N* individuals, each bearing a single chromosome containing 100 regions within which deleterious mutations occur at a total rate *U* per genome. The number of deleterious mutations within each region is tracked, incrementing the count for a region by one if it bears a mutation. Specifically, each generation, the total number of mutations is drawn from a Poisson distribution with mean *NU* and randomly assigned across the 100 regions of the entire population. During meiosis, recombination between regions is allowed, with a random number of crossovers drawn for each offspring, assuming a total map length of *L*; no recombination occurs within regions (otherwise there is no crossover interference). The fitness of an individual is computed as:

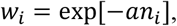

where *a* is the independent fitness effect of a mutation and *n*_*i*_ is the number of deleterious mutations carried by the individual. Both recombination rate and mutation rate are assumed to depend on the genotype at a single modifier locus, randomly placed in the genome. For each parameter combination, ten replicate burn-ins are conducted for 3*N* generations, and the populations are stored. For each burn-in population, the new modifier allele is introduced in a random region and tracked until it is lost or fixed, repeating the introduction process 5*N* times to obtain a mean fixation probability. We compared this fixation probability to the fixation probability for a new neutral mutation (1/*N*).

Simulation results confirm that modifier alleles that increase recombination are more likely to spread because they reduce selective interference among deleterious mutations (Keightley and Otto 2006). Modifier alleles that expand the genetic length of a chromosome from 0 Morgans (M) to 0.1 M (**Figure 3A**) or from 0.1 M to 0.2 M (**Figure 3B**) are favored. The advantage is much larger when there is little recombination initially (**Figure 3A**). In this scenario, a modifier allele that also increases the genomic deleterious mutation rate (*U*) by 5% or 10% is still favored but is less likely to spread than a modifier allele that does not elevate the mutation rate (**Figure 3A**). When the chromosome initially recombines, however, inducing a higher mutation rate effectively cancels the benefit accrued from more recombination, bringing the fixation probability of a modifier allele near or below that of a neutral allele (**Figure 3B**).

**Figure 3.**
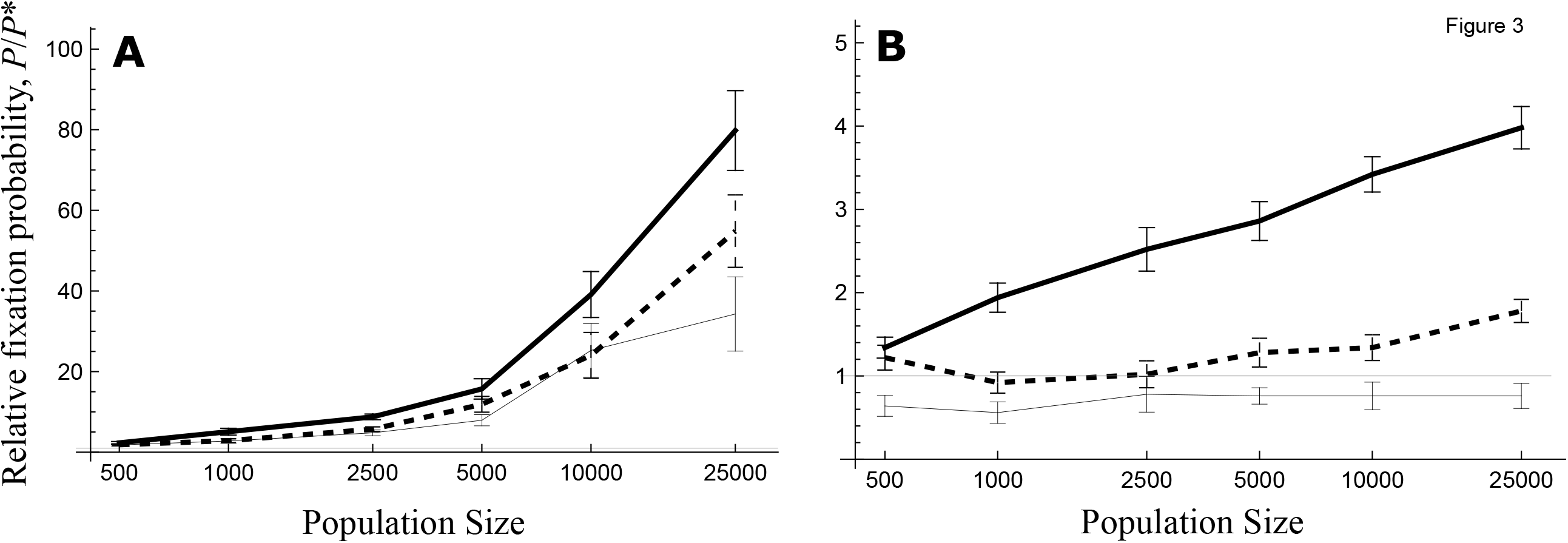
The probability of fixation for a modifier allele that controls recombination rate and mutation rate in finite populations with drift and selection. The fixation probability, *P*, of an additively acting modifier allele that increases the genetic length of a chromosome by 0.1 Morgans (M) in homozygotes, relative to the fixation probability of a neutral allele, *P*^*^, is shown as a function of population size *N* for cases of chromosomes with initial map lengths *L* = 0 M (A) and *L* = 0.1 M (B). The chromosomal mutation rate and fitness parameters are *U* = 1 and *a* = 0.01. Simulations model selective interference between deleterious mutations with no epistasis, a per mutation fitness parameter of *a* = 0.01, and a base chromosomal mutation rate of *U* = 1 (as in Figure 1a,b of Keightley and Otto 2006). Lines denote no mutagenic effect (thick black), 0.05 mutations added in new modifier homozygotes (dashed), 0.1 mutations added in new modifier homozygotes (thin), and neutral allele for comparison (horizontal gray). Means and standard error bars are calculated across ten replicate burn-ins.

The difference in relative fixation probability for a modifier allele with and without mutational effects grows with population size (**Figure 3**). These results demonstrate that recombination-associated mutation inhibits the evolution of recombination, even in finite populations without epistasis.

## Discussion

The process of forming crossovers elevates the mutation rate (Magni 1963; Strathern et al. 1995; Rattray et al. 2015; Halldorsson et al. 2019; Hinch et al. 2023). Our analysis demonstrates an important evolutionary consequence of this phenomenon. Even small increases in mutation narrow the zone in which higher recombination rates can evolve.

Recombination-associated mutation could impose upper limits on the evolution of recombination. The number of crossovers per chromosome falls within the range of one to three across a wide variety of species (Stapley et al. 2017; Otto and Payseur 2019). Although the lower bound likely reflects meiotic requirements to ensure proper segregation (Mather 1938; Hassold and Hunt 2001), determinants of the upper bound are less clear (Coop and Przeworski 2007). A simulation study found that the distribution of recombination rate within a human population is consistent with declines in fitness when the rate is high (Drury et al. 2023).

Our results suggest that evolutionary expansions of the genetic map are likely to be accomplished through those changes that simultaneously minimize associated mutagenesis. Some genome-wide association studies (GWAS) of recombination and some patterns of molecular evolution broadly match this prediction. Although candidate genes for variation in crossover number within populations (identified by GWAS) function at multiple steps in the pathway, there is an enrichment for genes that act after DSB generation, particularly those involved in the decision to repair DSBs as crossovers (Johnston 2024; Payseur 2025). Rapidly evolving recombination genes in mammals are functionally clustered around the crossover/non-crossover choice and the formation of the synaptonemal complex (Dapper and Payseur 2019), although signatures of positive selection are spread more evenly across the recombination pathway in birds (Szasz-Green et al. 2025). To the extent that these patterns point away from variants that increase recombination by adding DSBs, they are consistent with our theoretical findings.

At the same time, increasing DSBs could have additional fitness benefits that balance the cost of higher mutation. For example, meiotic DSBs aid chromosome homology searching and pairing in many species (Gray and Cohen 2006), including those with strong evidence for recombination-associated mutation (*e*.*g*. yeast and humans). Models that explicitly treat the evolution of DSBs could provide new insights into the evolution of recombination.

We have focused our analysis on the advantage of increasing recombination when there is synergistic epistasis against deleterious mutations, combined with simulations of selective interference in finite populations without epistasis. Other scenarios may more strongly favor increased recombination, including non-random mating (Roze and Lenormand 2005) and environments that fluctuate temporally or spatially (Barton 1995; Otto and Michalakis 1998; Lenormand and Otto 2000). While we assumed that increases in mutation rate are inherently disfavored (Karlin and McGregor 1974; Altenberg et al. 2017), modifier alleles that confer higher mutation rates along with more recombination could evolve more readily in a changing environment when adaptive mutations are possible.

Another source of mutation ignored by our analyses is GC-biased gene conversion, the preferential transmission of G/C alleles over A/T alleles in heterozygotes caused by a bias in recombination-associated DSB repair (Eyre-Walker 1999; Galtier et al. 2001). To the extent that GC-biased gene conversion leads to higher rates of deleterious mutation connected to recombination (Galtier et al. 2009; Riffis et al. 2025), our results suggests this process could constrain the evolution of recombination (though modeling of this scenario is warranted).

The cost of recombination-associated mutation also depends on genome architecture. To use existing theoretical frameworks, we restricted mutation and recombination to the same viability loci, assuming mutations in these genes are deleterious. Our results might overestimate the impact of recombination-associated mutation if DSBs occur disproportionately in non-functional regions of the genome. For example, *PRDM9* directs recombination initiation away from promoters and enhancers in mice (Brick et al. 2012). Elevated mutation could pose a stronger barrier to the evolution of recombination when such positioning is not possible, for example, in species with high gene density, such as yeast. As is the case for most parameters that determine how recombination rate evolves, the cost of error-prone DSB repair will vary among species and among genomic regions.

## Appendix

### Recursions

Our analysis follows changes in three-locus haplotype frequencies from one generation to the next that result from mutation, recombination, and selection. With two alleles at each of the three loci, there are eight possible haplotypes {*MAB, MAb, MaB, Mab, mAB, mAb, maB, mab*} whose frequencies among adult haploids are {*x*_1_, *x*_2_, *x*_3_, *x*_4_, *x*_5_, *x*_6_, *x*_7_, *x*_8_}. Recombination and mutation occur simultaneously during meiosis in diploids, according to the transmission Table S1. For example, a diploid *MAB/mAb* produces *MaB* offspring only if mutation occurs at the *A* locus and not at the *B* locus, accompanied by either no recombination or double recombination, occurring with frequency *u*_*Mm*_(1 − *u*_*Mm*_)((1 − *R*) (1 − *r*_*Mm*_)/2 + *R r*_*Mm*_/2). The frequency of each haploid offspring (*e*.*g*., *x*_3,*rm*_ for *MaB*) is then obtained by multiplying the frequency of each diploid parent by the chance of producing that offspring and summing over all diploids.

These haploids then undergo selection, leading to the next generation of haploid adults:

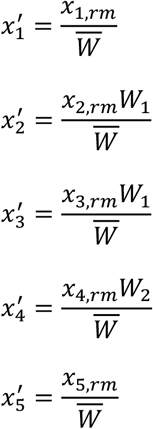

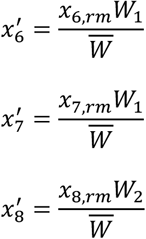

where 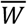 is the average fitness computed as the sum of haplotype fitnesses weighted by haplotype frequencies, *W*_1_ = 1 − *s*, and *W*_2_ = (1 − *s*)^2^ + *ε*.

### Equilibria

We first seek equilibria when the resident allele (*M*) is fixed in the population. To simplify the analysis, we assume that selection acts symmetrically, which reduces the system of four recursions involving *M* to two recursions (for *x*_1_ and *x*_2_), and we replace haplotype frequencies with expressions involving the frequency of deleterious alleles (*p* = *p*_*a*_ = *p*_*b*_ = 1 − (*x*_1_ + *x*_2_)) and the linkage disequilibrium between alleles (*D* = *D*_*AB*_). To find approximations for equilibrium values of *p* and *D*, we use a perturbation analysis assuming epistasis is weak (*ϵ* is small). We rewrite *p* and *D* as power functions of *ϵ*, insert them into the recursions, take Taylor series around *ϵ* = 0, and solve for the terms in zero order of *ϵ* and first order of *ϵ*. This procedure leads to equation (1) for the resident equilibrium in the main text.

### Stability to Introduction of Allele that Modifies both Mutation and Recombination

To determine the conditions under which an allele that modifies both mutation and recombination will spread through the population, we use a perturbation analysis. We approximate the leading eigenvalue (*λ*_*L*_) of the 4 x 4 Jacobian matrix involving the new modifier allele (*m*) evaluated at the resident equilibrium. We ask how small changes in *ϵ* or *d* perturb *λ*_*L*_ away from the boundary between stability and invasion (*λ*_*L*_ = 1). In the characteristic polynomial evaluated at the resident equilibrium, we replace *λ*_*L*_with 1 plus a small term of order *ϵ* and replace *d* with a small term of order *ϵ*. Assuming *ϵ* and *d* are small, taking Taylor series around *ϵ* = 0, and solving for the small term of order *ϵ* leads to expression (2) for *λ*_*L*_in the main text.

### Effects of Modifier of Mutation Rate in Two-Locus vs. Three-Locus Models

To understand how the effects of changing the mutation rate are expected to scale to a larger number of selected loci, we analyzed a simpler, two-locus model, in which a modifier locus controls the mutation rate at a single locus. The two-locus model features the same parameters and setup as the three-locus model, except the modifier does not affect recombination and viability is determined by a single locus. With two alleles at each locus, there are four haplotypes {*MA, Ma, mA*, and *ma*} with frequencies {*x*_1_, *x*_2_, *x*_3_, and *x*_4_}. Following mutation at the viability locus and recombination between the modifier and the viability locus, haplotype frequencies are:

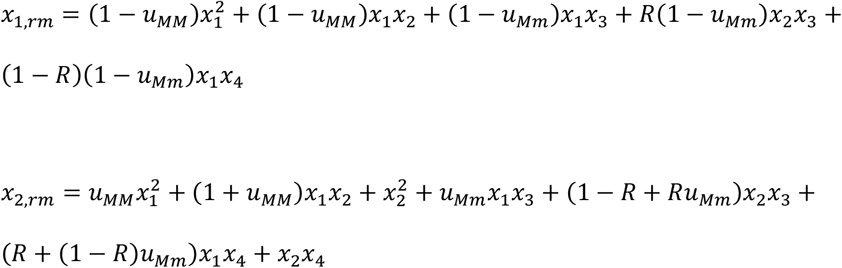

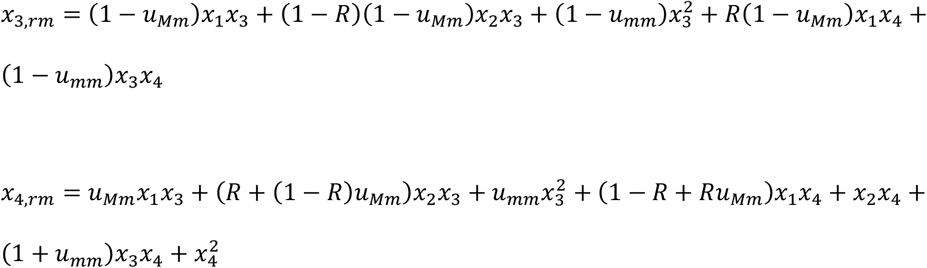

Following selection, haplotype frequencies in the next generation are:

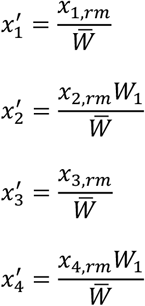

where 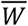 is average fitness computed as the sum of haplotype fitnesses weighted by haplotype frequencies, and *W*_1_ = 1 − *s*.

Solving for the equilibrium value of *x*_1_ with the resident allele (*M*) fixed (*x*_3_ = *x*_4_ = 0), leads to this expression:

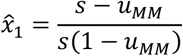

To determine the stability of this equilibrium, we construct the 2 x 2 Jacobian matrix for the two haplotype frequencies (*x*_3_, *x*_4_) involving the modifier allele that affects the mutation rate (*m*) and evaluate this matrix at the resident equilibrium. Solving for *λ*_*L*_ and again assuming the mutation effect of the modifier allele (*d*) is weak (of order *ϵ*), *λ*_*L*_is approximately:

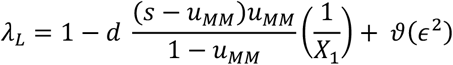

where *X*_1_ is again *X*[*R*] = *R*(1 − *s*) + *s* − *u*_*MM*_. Stability to invasion by the modifier allele is determined by the sign of *d*. When the modifier allele increases the mutation rate (*d* > 0), *λ*_*L*_ < 1, and it does not invade. When the modifier allele decreases the mutation rate (*d* < 0), *λ*_*L*_ > 1, and it spreads through the population.

To compare this result to the three-locus model, we sum the impact on a modifier from the viability locus *A* at distance *R*, given by the previous equation, and the impact on a modifier from the viability locus *B*, which lies at a distance *R*(1 − *r*_*Mm*_) + *r*_*Mm*_(1 − *R*) from the modifier (*i*.*e*., recombination must happen between intervals *M-A* or *M-B* but not both). Given that *X*[*R*(1 − *r*_*Mm*_) + *r*_*Mm*_(1 − *R*) ] was defined as *X*_2_, this sum returns the exact same coefficient for *d* as in equation (2). This result suggests that the effect of recombination-associated mutation on the spread of a recombination rate modifier scales in a nearly linear manner with the number of viability loci, *n* (proportional to 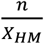, where *X*_*HM*_ is the harmonic mean value of *X*[*ρ*] over the affected loci).

## Acknowledgments

This research was supported by NIH R35GM139412 to BAP and NSERC RGPIN-2022-03726 to SPO. BAP thanks Linnea Sandell, Ailene MacPherson, Rob Unckless, and members of the Payseur lab for encouragement. We thank the reviewers for feedback that improved the manuscript. The authors declare no conflicts of interest.

## Author Contributions

BAP conceived the study. BAP and SPO developed and analyzed the model. BAP wrote the first draft of the manuscript and SPO revised it.

## Data and Code Availability

Code for analyses and simulations is available on GitHub at https://github.com/PayseurLabUWMadison/mutagenic_recombination_theory. Two Mathematica notebooks, “ThreeLocus_Manuscript.nb” and “TwoLocus_Manuscript.nb,” provide additional details on models and their analyses. These notebooks are also provided as pdf files. Code for simulations, modified from code written originally by Peter Keightley, is also provided.

